# A dominant role of cell death in limiting Chandipura virus propagation at cell-saturating high multiplicity of infection

**DOI:** 10.1101/2025.09.04.673995

**Authors:** Bhawna, Swapnava Basu, Syed Yusuf Mian, Sanchi Arora, Yashika Ratra, Kasturi Ganguly, Sachendra S. Bais, Manidipa Banerjee, Abhyudai Singh, Soumen Basak

**Author notes:** Harvard T.H. Chan School of Public Health, USA. Department of Medicine, Washington University School of Medicine, Saint Louis, MO 63130, USA. these authors contributed equally to this work (Bhawna, Swapnava Basu).

## Abstract

Viruses transit from a low to high multiplicity of infection (MOI) regime in infected tissues. Type-1 interferons (IFNs) enforce a cellular state refractory to virus multiplication, while the death of infected cells eliminates viral replicative niche. Here, we investigated how these two innate antiviral mechanisms cooperate at various MOIs upon cell infection by Chandipura virus (CHPV), a cytopathic RNA virus implicated in several outbreaks of acute encephalitis in India. We found that as expected, a gradual increase in the input MOI from 0.02 to 2 led to a proportionate surge in the viral yield. Surprisingly, a further rise to MOI 20 caused a reduction in the progeny titer. Our mathematical modeling together with *ex vivo* infection studies involving mutant cells suggested that cell death - more so than virus-induced type-1 IFNs - restricted CHPV propagation at cell-saturating high MOIs, leading to a net fall in the yield at MOI 20. We argue that the distinct involvement of innate immune pathways at varied MOIs imparts robust cellular defense against cytopathic viruses.

**Significance sentence:** Cell death - more so than type-1 interferons - limits Chandipura virus propagation at cell-saturating high multiplicity of infection.

**Highlights:** # Type-1 IFNs and cell death cooperate at varied MOI in limiting CHPV multiplication.

\# At sub-saturating low MOI, type-1 IFNs play a dominant role in controlling CHPV propagation.

\# At cell-saturating high MOI, cell death determines the progeny yield.

## Introduction

The intimately connected global civilization of the present era is also extremely vulnerable to epidemic threats. In particular, Chandipura virus (CHPV) has been linked to several outbreaks of acute encephalitis in India, including the most recent one in 2024 (Brisse & Ly, 2024; Devi, 2024; Mallick et al., 2025; Sapkal et al., 2018). Previous epidemiological studies captured a startling case fatality rate of 55% to 75% in CHPV outbreaks, with accompanying dysfunctions in the central nervous system, among children below fifteen years of age (Sudeep et al., 2016). An improved understanding of molecular and cellular events driving pathogenesis during the course of CHPV infection may have a profound bearing on human health, particularly in the Indian subcontinent.

CHPV belongs to the Rhabdoviridae family and Vesiculovirus genera and possesses an enveloped negative-sense RNA genome (Basak et al., 2007; Pandey & Singh, 2025). A recent study implicated host-derived α-2-macroglobulin in viral entry into target cells, although the exact identity of the cellular receptor for CHPV remains uncertain (Dey et al., 2024). Nevertheless, a series of prior investigations aptly defined the role of CHPV-encoded Phosphoprotein and Nucleocapsid protein in regulating the viral transcription-replication cycle in infected cells (Basak et al., 2003; Majumdar et al., 2004; Mondal et al., 2012; Raha et al., 2000). Similarly, cellular factors that collaborate with viral Matrix protein to promote CHPV budding from infected cells have been identified (Rajasekharan et al., 2015). Accordingly, the emphasis of CHPV research gradually shifted towards understanding host responses.

Type-1 IFNs and cell death constitute two principal arms of the innate antiviral cellular defense. Viral infection stimulates the nuclear translocation of RelA NF-κB heterodimers from their cytoplasmic IκBα-inhibited latent complexes and also activates Interferon regulatory factor 3 (IRF3). IRF3, in collaboration with RelA NF-κB, transcriptionally upregulates type-1 IFNs, including IFNβ. In autocrine and paracrine loops, type-1 IFNs secreted from infected cells then signal through the cognate IFN receptor (IFNR) – this triggers hundreds of interferon-stimulated genes (ISGs) (McNab et al., 2015). ISGs enforce an antiviral cellular state refractory for viral multiplication. Accordingly, genetic disruption of type-1 IFN signaling in *Ifnar1*^*-/-*^ cells, which lack functional IFNR, was shown to enhance the multiplication of a number of RNA viruses, including CHPV (Abraham et al., 2010; Bais et al., 2019; Ratra et al., 2022).

Infection-inflicted cell death also limits viral multiplication, albeit by eliminating the viral replicative niche (Jorgensen et al., 2017; Orzalli & Kagan, 2017). CHPV represents a cytopathic RNA virus. CHPV infection is associated with apoptosis driven by FAS-associated death domain (FADD)-mediated caspase 8 activation and also necroptosis involving receptor-interacting serine-threonine protein kinase 3 (RIPK3)-directed phosphorylation of mixed lineage kinase domain-like (MLKL) (Bais et al., 2019; Ghosh et al., 2013). Not surprisingly, viruses tend to counteract the death of infected cells to ensure their continued growth. In fact, virus-activated RelA NF-κB also drives the expression of pro-survival genes, which prolong the life of infected cells. Correspondingly, a genetic deficiency of RelA NF-κB in mouse embryonic fibroblasts (MEFs) was shown to escalate the activation of apoptotic caspases and MLKL upon CHPV infection, leading to exacerbated cell death and a decrease in the viral yield (Bais et al., 2019).

Despite an in-depth understanding of the individual type-1 IFN and cell death pathways, how these two arms mutually cooperate remains less clear. In particular, previous studies documented that the effective multiplicity of infection (MOI) rises substantially within the tissue during the course of infection by animal and also plant viruses (Gutiérrez et al., 2010; Josefsson et al., 2011; Rüdiger et al., 2019). Consistently, low MOI cell infection using Influenza A virus revealed a gradual transition from low to high MOI settings (Rüdiger et al., 2019). However, a plausible impact of altered input MOI on CHPV-induced type-1 IFN response and cell death, and also progeny yield, has not been examined.

Our combined experimental and mathematical studies identified a MOI-dependent collaboration between type-1 IFNs and cell death processes. We found that the type-1 IFN pathway was an important determinant of CHPV yield in the sub-saturating low MOI infection regime, whereas cell death represented a dominant antiviral mechanism during cell-saturating high MOI infections. In sum, our work suggests that distinct engagement of innate antiviral processes at varied MOIs instills robust cellular defense against cytopathic viruses.

## Results

### A paradoxical relationship between viral input and yield at cell-saturating MOIs

To understand how the quantity of initial infecting particles impacts virus yield, we infected MEFs with CHPV at 0.02, 0.2, 2, and 20 MOI and determined the progeny titer in the culture supernatant by standard plaque assay (Fig 1a). As expected, we recorded a proportionate rise in the titer, measured at 24hr post-infection, with a gradual increase in the input MOI from 0.02 to 2 (Fig 1b). Unexpectedly, when the MOI was further raised from 2 to 20, we noticed a 2.2-fold reduction in the progeny yield. To ensure that our observation was not specific for a particular time point, we carried out a series of measurements at 12hr, 24hr, and 36hr post-infection, comparing CHPV yield at cell-saturating 2 and 20 MOIs. CHPV infection at MOI 20 produced, if anything, subtly more progeny virus at the early 12hr time point (Fig 1c). We noticed a rise in the progeny yield in the cell culture supernatant with time for both MOIs. This increase was less remarkable for cell infection at MOI 20 than MOI 2, culminating to an overall deficit in the progeny titer at 24hr and 36hr time points for the MOI 20 regime.

**Fig 1.**
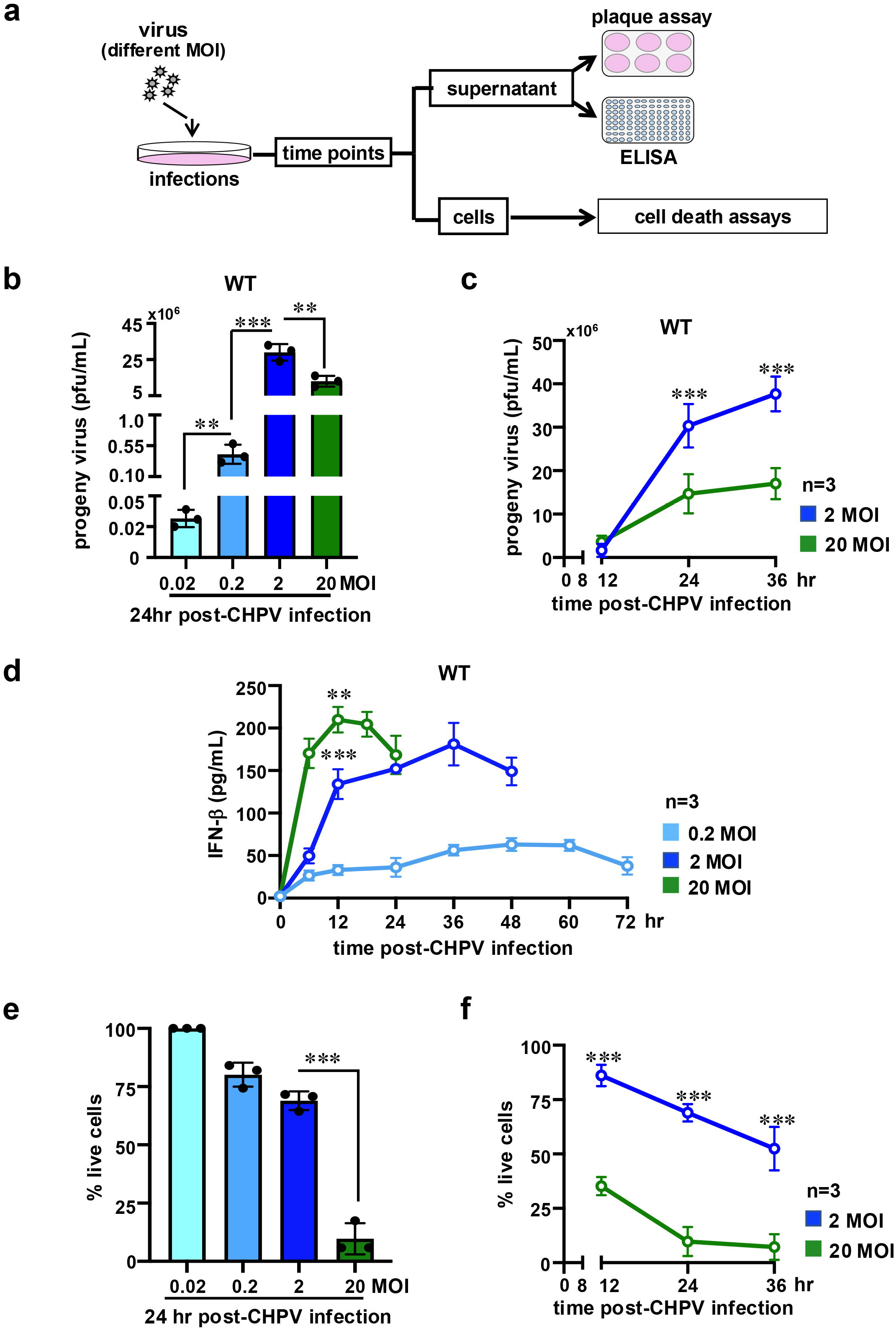
Comparing CHPV propagation in cultured cells infected at different MOIs. *(a)* Schema summarizing cell-infection experiments performed in this study. *(b)* Barplot revealing progeny virus titer in the culture supernatant of MEFs at 24hr post-infection with CHPV at the indicated input MOI. *(c)* Similarly, progeny virus yield was measured at 12hr and 36hr post-infection for cells infected at MOI 2 and 20. *(d)* ELISA showing accumulation of IFNβ in a time course in the culture supernatant of cells infected with CHPV at the indicated MOI. *(e)* CHPV-mediated cell death was measured at 24hr post-infection by crystal violet staining. The abundance of viable cells at various MOI was determined relative to corresponding uninfected MEFs and presented as bargrams. *(f)* Infection-induced cell death was similarly measured at 12hr and 36hr post-infection. Data represent the means of three biological replicates ± SEM. The statistical significance was determined using two-way ANOVA test. \**P* ≤ 0.05; \*\**P* ≤ 0.01; \*\*\**P* ≤ 0.001.

Next, we charted the innate antiviral cellular processes at various MOIs. We determined the abundance of IFNβ in the culture supernatant of infected cells by ELISA. Our analyses revealed that an increase in MOI both enhanced and accelerated IFNβ production. In particular, MOI 2 and MOI 20 infections produced an average peak IFNβ concentration of 181.07 pg/ml and 204.44 pg/ml in the culture supernatant at 36hr and 12hr post-infection, respectively (Fig 1d). However, we noticed an early drop in the IFNβ production at 18h post-infection in the MOI 20 regime. Using crystal violet staining, we then captured cell death as a function of input MOI. We found that increasing MOI caused a steady decline in the cell viability at 24hr post-infection, leading to 80% and 69% live cells at MOI 0.2 and MOI 2, respectively (Fig 1e). Curiously, cell death was more strikingly escalated at MOI 20, leaving only a residual 9.7% live cells. Our time course analyses substantiated a significantly low live cell count at MOI 20 compared to MOI 2 as early as 12hr (Fig 1f). We conclude that increasing input MOI at the sub-saturating regime augments CHPV yield while also intensifying type-1 IFN and cell death responses. At saturating MOIs, however, a further rise to high MOI values paradoxically plunges the progeny yield.

### Mathematical studies suggest distinct engagements of type-1 IFNs and cell death in limiting viral yield at varied MOI

Mathematical modelling provides valuable insights into the dynamic interplay between viral multiplication and host responses (Boianelli et al., 2015; Van Eyndhoven et al., 2021; Wodarz & Thomsen, 2005). In particular, previous ordinary differential equation (ODE)-based studies suggested that while IRFs tune subtype-specific type-1 IFN responses to Influenza A virus, the extent of cell death could be a more accurate indicator of the severity of infections (Weaver & Shoemaker, 2020). Mathematical analyses also revealed that the duration, and not only the amplitude, of type-1 IFN feedback is crucial for eliminating infectious VSV particles, particularly in the context of heightened viral replication rates (Tan et al., 2012). Similarly, combined single-cell studies and mathematical modeling experiments elucidated that an epigenetic-driven mechanism enforces transient heritability among the fraction of cells producing type-1 IFNs in a population (Van Eyndhoven et al., 2023). Our own mathematical studies identified that RelA NF-κB may promote CHPV yield by limiting infection-inflicted cell death (Bais et al., 2019). Although these investigations underscored the role of cell death and type-1 IFNs in viral growth, how input MOI impacts these dynamic relationships has not been mathematically dissected.

Because our experiments revealed a discrepant decrease in the progeny titer upon increasing the input MOI from 2 to 20, we set out to mathematically dissect CHPV growth. To this end, we developed a three-state ODE-based model to describe CHPV multiplication in cultured cells infected at saturating MOIs (Fig 2a). We assumed that all cells were infected at the start of the experiment. After a time delay of *t1*_*delay*_, infected cells produced IFNs involving a rate constant *IFN*_*pdn*_ and progeny viruses were generated involving a rate constant of *Virus*_*pdn*_. We also considered a basal IFN level of 2.5pg/ml in our model, as experimentally determined in WT cells (Fig 1d, Table 1a). After an additional delay of *t2*_*delay*_, cell death occurred involving a rate constant of *Cell*_*death*_. *IFN*_*inh*_ was used as the rate constant for IFN-mediated inhibition of virus production by infected cells. The degradation rate of IFNs and the rate for inactivation of viruses were represented using *IFN*_*deg*_ and *Virus*_*deg*_, respectively. Subsequently, we fitted our model to experimental time course data (Fig 2b, Supplementary Information). From the fitting exercise, we then deduced *t1*_*delay*_ and *t2*_*delay*_ values to be 2.5hr and 3.5hr, respectively, regardless of the input MOI (Materials and Methods, Supplementary Information). Similarly, we estimated the values for *IFN*_*deg*_, *Virus*_*deg*_ and *IFN*_*inh*_ to be 7×10^-3^ hr^-1^, 1.0×10^-5^ hr^-1^ and 1.0×10^-5^ hr^-1^, respectively. The mean extracted values for *Virus*_*pdn*_, *IFN*_*pdn*_ and *Cell*_*death*_ at MOI 2 were 13×10^-4^ cell^-1^hr^-1^, 1.9 pg mL^-1^hr^-1^ and 0.024 hr^-1^, respectively. We further determined that while leading to a rather insignificant alteration in *Virus*_*pdn*_, a change in the input MOI from 2 to 20 caused a 2.3-fold rise in *IFN*_*pdn*_ (Fig 2c, Table 1b). Interestingly, our mathematical formalism projected a marked 8-fold increase in *Cell*_*death*_ from 0.02 hr^-1^ to 0.16 hr^-1^ when MOI was changed from 2 to 20 (Fig 2c, Table 1b). Taken together, a rise in the input MOI prompted heightened type-1 IFN response and aggravated cell death.

**Table 1a:** A chart describing the list of rate parameters and their corresponding values determined by fitting experimental data into the mathematical model.

| Parameter | 2 MOI |  |  | 20 MOI |  |  |
| --- | --- | --- | --- | --- | --- | --- |
|  | WT | *Ifnar1*<sup>-/-</sup> | *Nfkb1a*<sup>-/-</sup> | WT | *Ifnar1*<sup>-/-</sup> | *Nfkb1a*<sup>-/-</sup> |
| IFN(0) pg/ml (basal) | 2.5 | 2.5 | 5.25 | 2.5 | 2.5 | 5.25 |
| *t*<sub>1delay</sub> (hr) | 2.5 | 2.5 | 2.5 | 2.5 | 2.5 | 2.5 |
| *t*<sub>2delay</sub> (hr) | 3.5 | 3.5 | 3.5 | 3.5 | 3.5 | 3.5 |
| *Virus*<sub>pdn</sub> (pfu cell<sup>-1</sup> hr<sup>-1</sup>) | (13.6±4.7)*10<sup>4</sup> | (35.2±9.3)*10<sup>4</sup> | (0.17±0.06)*10<sup>4</sup> | (12.3±4.1)*10<sup>4</sup> | (42.7±11.53)*10<sup>4</sup> | (1.6±1.4)*10<sup>4</sup> |
| *IFN*<sub>pdn</sub> (pg mL<sup>-1</sup> hr<sup>-1</sup>) | 1.95±0.07 | 0.77±0.2 | 0.28±0.05 | 4.43±1.2 | 2.68±0.79 | 0.69±0.06 |
| *Cell*<sub>death</sub> (hr<sup>-1</sup>) | 0.02±0.006 | 0.06±0.015 | 0.01±0.007 | 0.16±0.04 | 0.38±0.13 | 0.04±0.01 |
| *IFN*<sub>deg</sub> (hr<sup>-1</sup>) | 7*10<sup>-3</sup> | 7*10<sup>-3</sup> | 7*10<sup>-3</sup> | 7*10<sup>-3</sup> | 7*10<sup>-3</sup> | 7*10<sup>-3</sup> |
| *Virus*<sub>deg</sub> (hr<sup>-1</sup>) | 10<sup>-5</sup> | 10<sup>-5</sup> | 10<sup>-5</sup> | 10<sup>-5</sup> | 10<sup>-5</sup> | 10<sup>-5</sup> |
| *IFN*<sub>inh</sub> (mL pg<sup>-1</sup>) | 10<sup>-5</sup> | 0 | 10<sup>-5</sup> | 10<sup>-5</sup> | 0 | 10<sup>-5</sup> |
| *C*<sub>basal</sub><br>[fraction of alive cells regardless of infection] | 0.1 | 0.1 | 0.1 | 0.1 | 0.1 | 0.1 |

**Table 1b:**
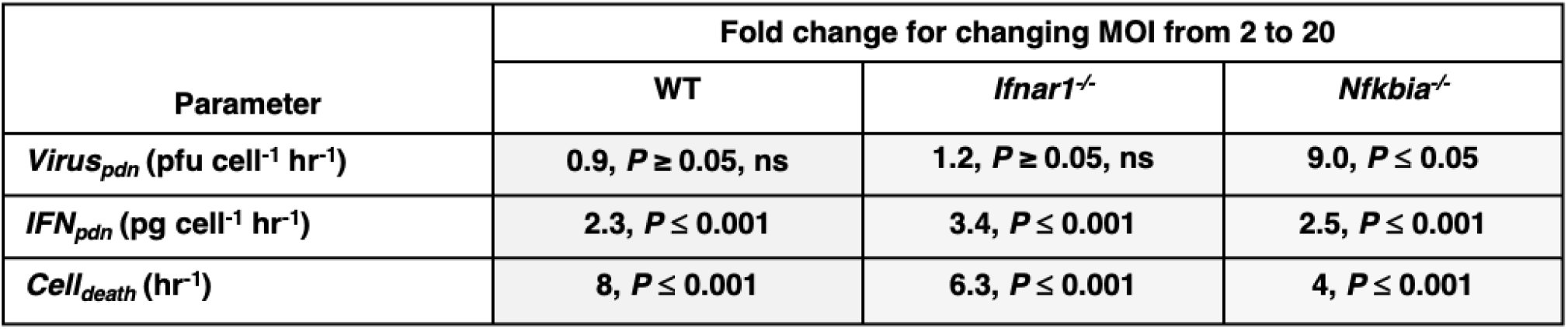
Comparing genotypes for MOI-dependent changes in the key rate parameters.

| Parameter | Fold change for changing MOI from 2 to 20 |  |  |
| --- | --- | --- | --- |
|  | WT | *Ifnar1*<sup>-/-</sup> | *Nfkb1a*<sup>-/-</sup> |
| *Virus*<sub>pdn</sub> (pfu cell<sup>-1</sup> hr<sup>-1</sup>) | 0.9, *P* ≥ 0.05, ns | 1.2, *P* ≥ 0.05, ns | 9.0, *P* ≤ 0.05 |
| *IFN*<sub>pdn</sub> (pg cell<sup>-1</sup> hr<sup>-1</sup>) | 2.3, *P* ≤ 0.001 | 3.4, *P* ≤ 0.001 | 2.5, *P* ≤ 0.001 |
| *Cell*<sub>death</sub> (hr<sup>-1</sup>) | 8, *P* ≤ 0.001 | 6.3, *P* ≤ 0.001 | 4, *P* ≤ 0.001 |

**Fig 2.**
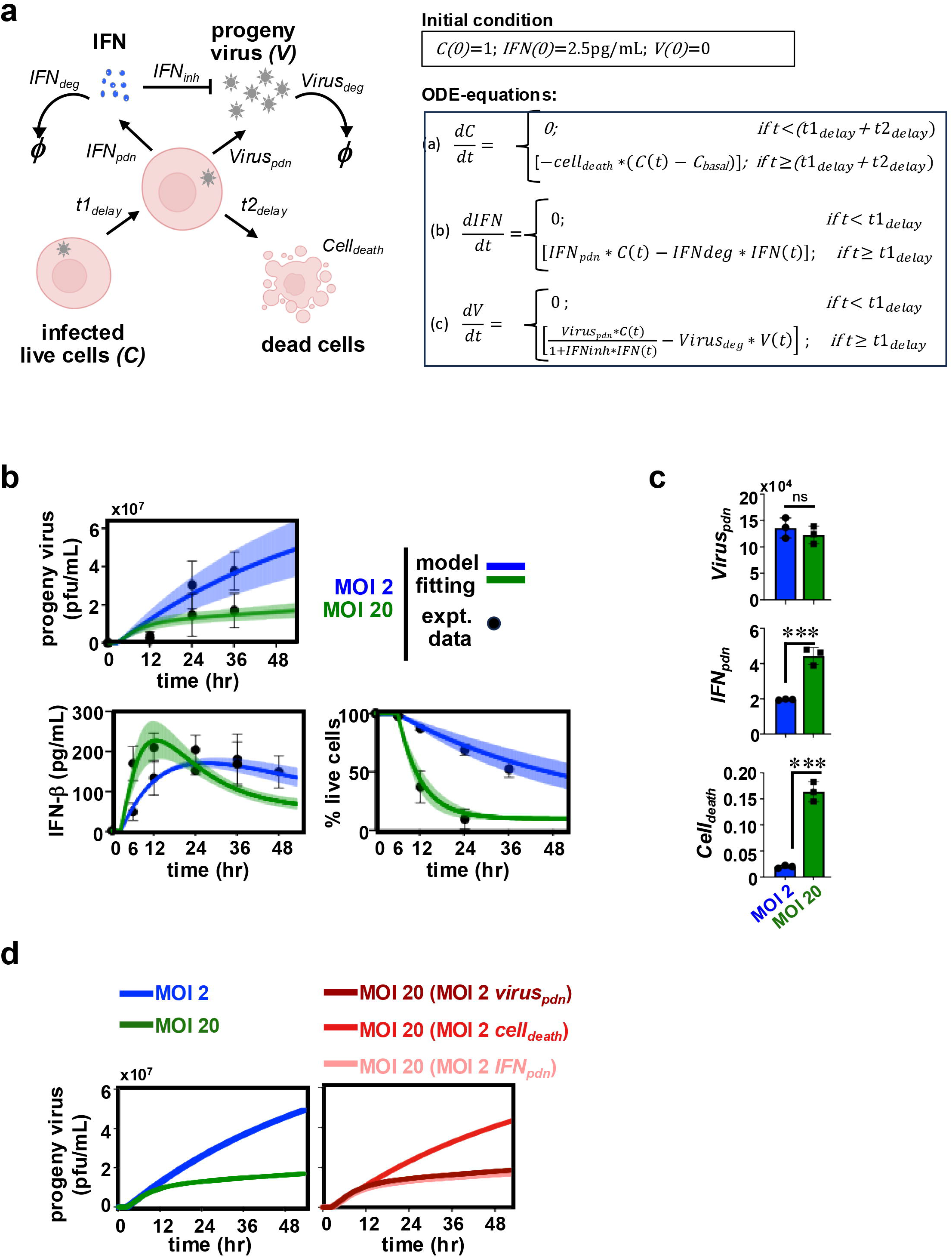
Mathematically probing CHPV multiplication and host responses at cell-saturating MOIs. *(a)* A cartoon depicting key processes involved in CHPV perpetuation at cell-saturating MOIs that were considered for mathematical analyses. Kinetic rate parameters associated with each of the reactions have been indicated. Bottom, a set of mathematical equations used for describing the relationship between no. of infected live cells, IFN concentration and progeny virus titer has been indicated. *(b)* Fitting mathematical equations with experimental time course data related to progeny virus titer, cell-produced IFNβ levels, and infection-inflicted cell death observed at MOI 2 and MOI 20. Experimental data points derived from biological replicates are indicated in black circles, and solid lines represent the fitted mean values for viral titre, IFN-β levels, or fraction of live cells. Shaded regions around the solid lines indicate the corresponding confidence intervals. *(c)* Bar chart comparing the values of *Cell*_*death*_, *IFN*_*pdn*_, and *Virus*_*pdn*_ at MOI 2 and MOI 20 extracted from fitting exercise. Values of these rate parameters were independently determined using data from experimental replicates. Unpaired t-test was performed to determine the statistical significance. *(d)* Simulating virus yield as a function of time in the MOI 20 regime using the *IFN*_*pdn*_ or *Cell*_*death*_ rate constant values linked to MOI 2. Data represent the means of three biological replicates ± SEM. \**P* ≤ 0.05; \*\**P* ≤ 0.01; \*\*\**P* ≤ 0.001; ns ∼ not significant, ≥ 0.05.

To understand how these kinetic rates may have influenced the viral yield at MOI 20, we charted virus generation over time in our mathematical model, albeit substituting MOI 20 rate values with those for MOI 2 (Fig 2d). Our mathematical analyses revealed that neither MOI 2-associated *Virus*_*pdn*_ nor MOI 2-associated *IFN*_*pdn*_ restored the virus titer in the MOI 20 regime. However, replacing the MOI 20 value with that of MOI 2 for *Cell*_*death*_ alone almost entirely rescued the yield at MOI 20, leading to a progeny titer comparable to the MOI 2 regime (Fig 2d). Our mathematical studies argue that escalating cell death, more so than elevated type-1 IFN production, restricts viral yield upon increasing input MOI in the cell-saturating MOI regime.

### Type-1 IFN deficiency fails to rescue the progeny yield at very high MOI

To experimentally verify the predictions from our mathematical analyses, we examined CHPV infections in *Ifnar1*^*-/-*^ cells, which lack functional type-1 IFN signaling. As such, type-1 IFN deficiency boosted viral yield, leading to a ∼170-fold enhancement in the progeny titer at 24hr post-infection in *Ifnar1*^*-/-*^ cells in the MOI 0.2 regime (Fig 3a). Notably, this IFN phenotype was considerably less profound at saturating MOIs, resulting in merely 2-fold and 2.8-fold differences in the viral titer between WT and *Ifnar1*^*-/-*^ cells at MOI 2 and MOI 20, respectively. We also noted that raising input MOI from 2 to 20 caused a reduction in the progeny yield in *Ifnar1*^*-/-*^ cells similar to those observed in WT MEFs. Likewise, increasing MOI led to a rise in the IFNβ level in the culture supernatant of *Ifnar1*^*-/-*^ cells, reaching a peak concentration of 106.5 pg/ml at 12hr post-infection in the MOI 20 regime (Fig 3b). However, the peak value for IFNβ was somewhat lower in *Ifnar1*^*-/-*^ MEFs than WT cells for a range of MOIs (compare Fig 1d and Fig 3b). *Ifnar1*^*-/-*^ cells also displayed slightly diminished viability as opposed to WT MEFs at MOI 0.2 that otherwise showed MOI-dependent deterioration, sparing less than 9% lives cells at 24hr post-infection in the MOI 20 regime (Fig 3c, Fig 3d).

**Fig 3.**
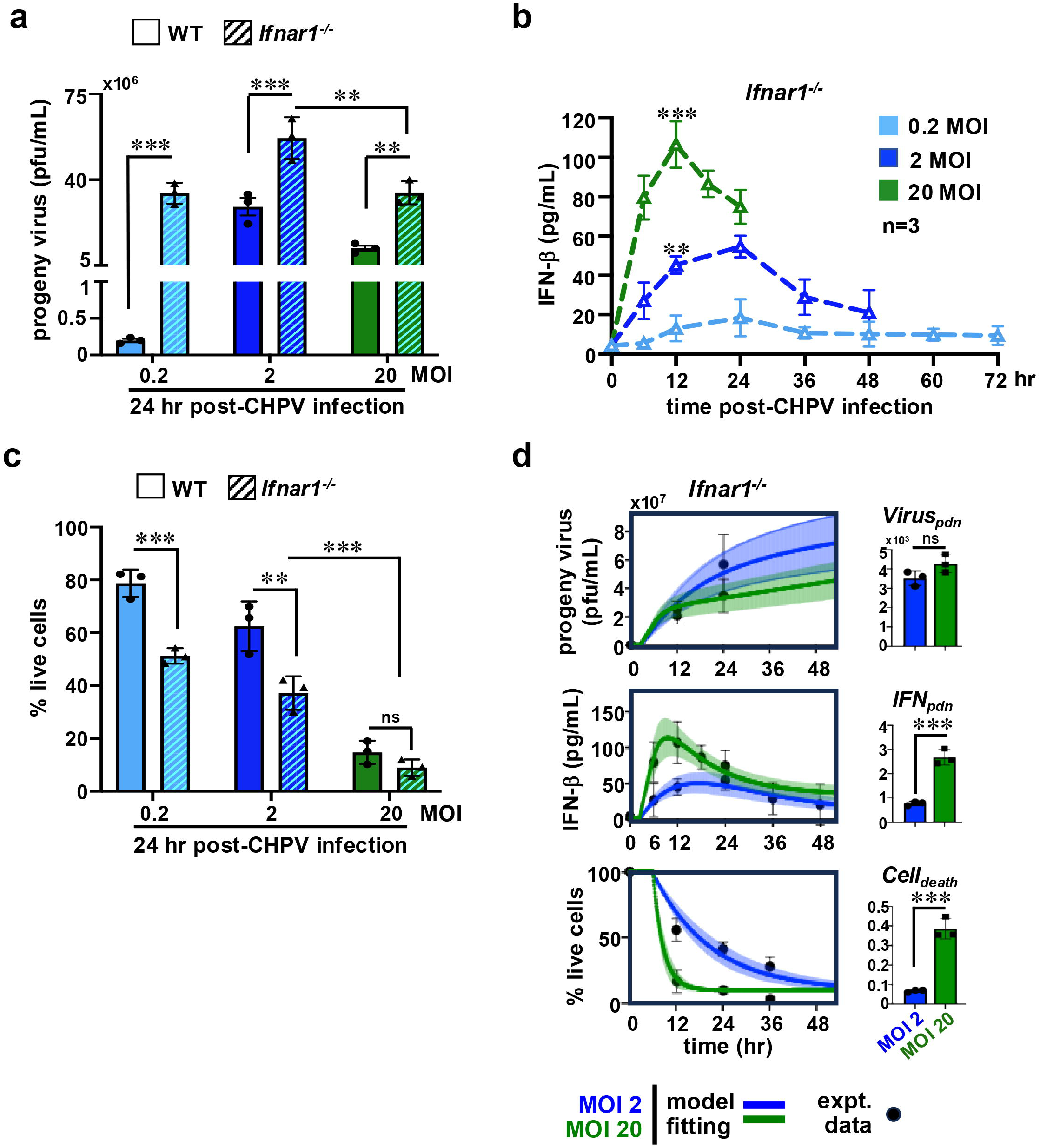
Charting CHPV growth in *Ifnar1*^*-/-*^ MEFs infected at different MOI. *(a)* Barplot comparing virus yields in WT and *Ifnar1*^*-/-*^ MEFs determined at 24hr post-infection at the indicated MOIs. *(b)* ELISA revealing the abundance of IFNβ in the culture supernatant of *Ifnar1*^*-/-*^ cells infected with CHPV at the indicated MOIs. *(c)* Barplot showing cell death in WT and *Ifnar1*^*-/-*^ MEFs at 24hr post-infection with CHPV at the indicated MOIs. *(d)* Line plot depicting fitting of mathematical equations with experimental time course data on CHPV propagation in *Ifnar1*^*-/-*^ cells (left). Extracted values for various rate parameters are indicated in the accompanying bar chart (right). Data represent the means of three biological replicates ± SEM. The statistical significance was determined using two-way ANOVA test for Fig 3a-Fig 3c and using unpaired t-test for Fig 3d. \**P* ≤ 0.05; \*\**P* ≤ 0.01; \*\*\**P* ≤ 0.001; ns ∼ not significant, ≥ 0.05.

We then fitted experimental time course data from *Ifnar1*^*-/-*^ cells infected at saturating MOIs to our mathematical model and extracted various rate parameters for understanding IFN-mediated virus control (Fig 3d, Table 1a). Because these cells lacked type-1 IFN signaling, *IFN*_*inh*_, the rate constant representing IFN-mediated inhibition of virus production, was set to zero. Otherwise, the values for *t1*_*delay*_, *t2*_*delay*_, *IFN*_*deg*_, and *Virus*_*deg*_ were deemed to be identical to those for WT. Our mathematical analyses disclosed that type 1 IFN deficiency led to a 2.6-fold to 3.5-fold increase in *Virus*_*pdn*_ and another 2.3 to 3.0-fold increase in *Cell*_*death*_, while resulting in a modest 1.6-fold to 2.5-fold reduction in *IFN*_*pdn*_ in these MOIs (Table 1a). Akin to WT cells, changing MOI from 2 to 20 within the *Ifnar1*^*-/-*^ settings produced a negligible change in *Virus*_*pdn*_, a moderate 3.4-fold increase in *IFN*_*pdn*_, and a prominent 6.3-fold rise in *Cell*_*death*_ (Table 1b). We infer that type-1 IFNs play only a minor role in MOI-dependent virus control, and in infection-induced cell death, at cell-saturating MOIs.

### Resilience to cell death improves the progeny CHPV yield at very high input MOI

Next, we asked if restraining virus-induced cell death bettered the progeny yield at high MOI in our experiments, as prescribed by our mathematical model. In addition to immune genes, RelA NF-κB dimers promote the expression of pro-survival factors, including cIAPs, cFLIP, Bcl2 and Bcl-XL (Gilmore, 2006). Our previous investigation demonstrated that the genetic absence of RelA aggravates cell death upon CHPV infection (Bais et al., 2019). In resting cells, RelA NF-κB factors are sequestered in the cytoplasm in a latent form by the inhibitory IκB proteins, the major isoform being *Nfkbia*-encoded IκBα (Mulero et al., 2019). It was shown that disruption of IκBα-mediated inhibition promotes basally elevated NF-κB activity in *Nfkbia*^*-/-*^ cells (Beg et al., 1995) and that constitutive NF-κB activity confers protection against death-inducing stimuli (Guo et al., 2024). Accordingly, we set out to examine *Nfkbia*^*-/-*^ MEFs to assess the role of cell death in CHPV propagation.

We found that IκBα deficiency indeed improved the cell viability, particularly more discernably at cell-saturating MOIs (Fig 4a). Enhanced viability of *Nfkbia*^*-/-*^ cells at MOI 20 led to a net 3.6-fold increase in the live cell count at 24hr post-infection compared to those in WT MEFs. Furthermore, *Nfkbia*^*-/-*^ cells maintained ∼2.2-fold excess IFNβ in the culture supernatant basally, while generating less IFNβ upon infection for a range of MOIs (Table 1a, also compare Fig 1d and Fig 4b). Although increasing MOI elevated IFNβ levels, *Nfkbia*^*-/-*^ cells did not display early termination of IFNβ production at MOI 20, as observed in other genotypes. These knockouts also produced substantially fewer progeny virus particles at varied MOIs (Fig 4c). Importantly, a lack of IκBα arrested the decline in the virus yield upon raising the input MOI in the cell-saturating regime, instead leading to an overall 9.5-fold enhancement in the progeny titer at MOI 20 compared to MOI 2 (Fig 4c).

**Fig 4.**
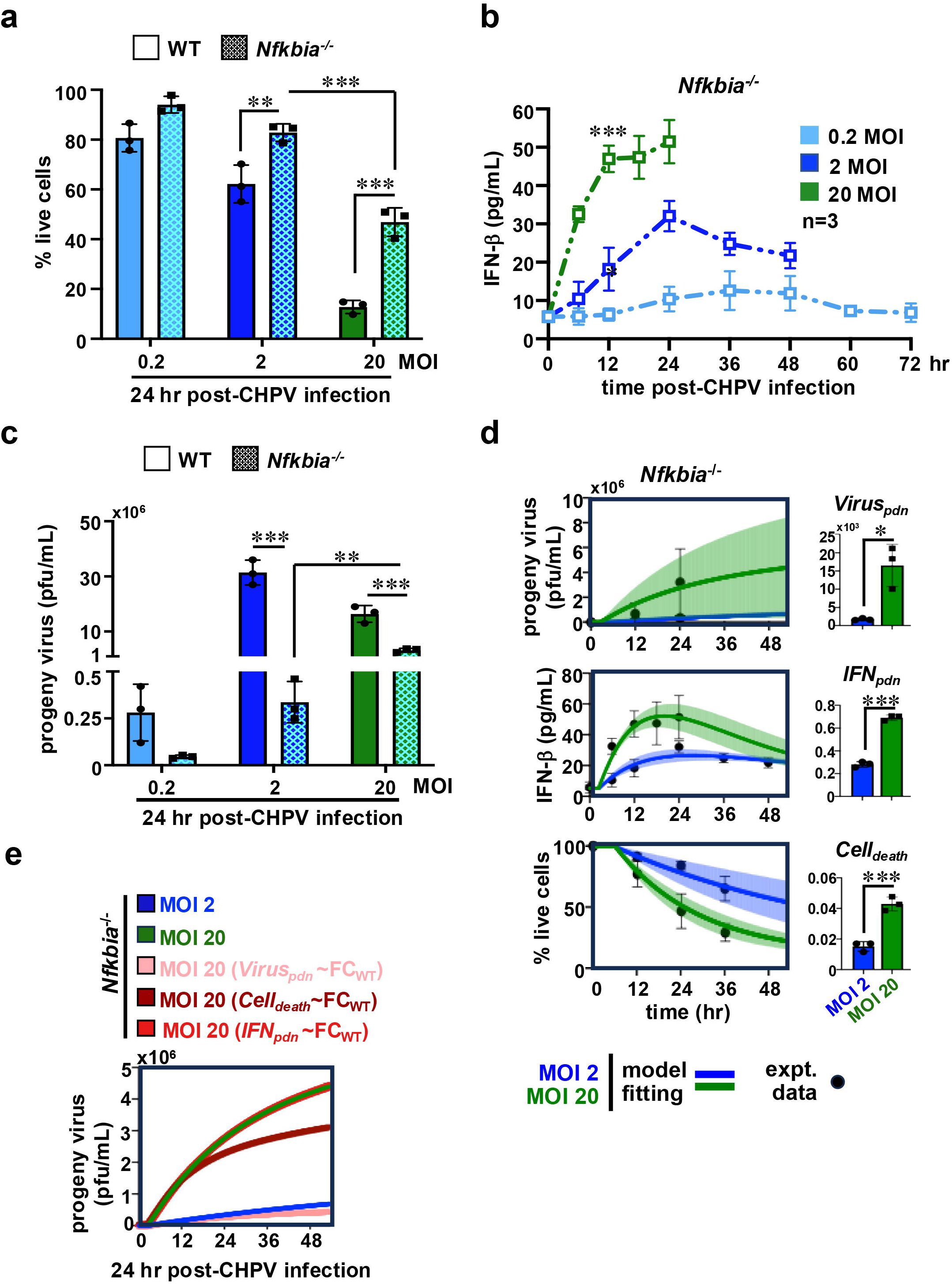
Studying CHPV propagation in *Nfkbia*^*-/-*^ MEFs infected at different MOI. *(a, b, c)* Comparing WT and *Nfkbia*^*-/-*^ MEFs infected at various MOIs for cell death (*a*), IFNβ production (*b*), and CHPV yield (*c*). *(d)* Fitting experimental time course data on CHPV propagation in *Nfkbia*^*-/-*^ cells in our mathematical model (left). Extracted values for various rate parameters are also indicated (right). *(e)* Line plot charting the simulated progeny virus titer estimated in the *Nfkbia*^*-/-*^ settings at MOI 20 using altered *Virus*_*pdn*_, *Cell*_*death*_, and *IFN*_*pdn*_ values that preserved the quantum of changes in these rates upon raising MOI from 2 to 20 to those determined in WT cells. Data represent the means of three biological replicates ± SEM. The statistical significance was determined using two-way ANOVA test for Fig 4a-Fig 4c and using unpaired t-test for Fig 4d. \**P* ≤ 0.05; \*\**P* ≤ 0.01; \*\*\**P* ≤ 0.001.

Fitting *Nfkbia*^*-/-*^ cell data obtained at saturating MOIs to our mathematical model, we then deduced the relevant rate parameters (Fig 4d, Table 1a). Our mathematical approach disclosed that IκBα deficiency led to a substantial 76.5-fold and 8.1-fold drop in *Virus*_*pdn*_ at MOI 2 and MOI 20, respectively, and an approximately 6.6-fold reduction in *IFN*_*pdn*_. Compared to WT, *Nfkbia*^*-/-*^ cells also displayed a 2-fold decrease in *Cell*_*death*_ at MOI 2, and a more prominent 4-fold fall at MOI 20. We then focused on probing how MOI changes from 2 to 20 altered these rates within the *Nfkbia*^*-/-*^ settings. Despite an overall decline in the virus yield compared to WT MEFs, we captured a marked 9-fold elevation in *Virus*_*pdn*_ in *Nfkbia*^*-/-*^ cells when MOI was raised to 20 (Table 1b). Notably, an equivalent increase in MOI led to a rather insignificant change in *Virus*_*pdn*_ in WT cells. While MOI-dependent changes in *IFN*_*pdn*_ were analogous between WT and *Nfkbia*^*-/-*^ MEFs, IκBα deficiency restricted the rise in *Cell*_*death*_ upon increasing MOI to 4-fold compared to 8-fold in WT cells (Table 1b). We then altered the MOI 20 values for *Virus*_*pdn*_, *Cell*_*death*_, and *IFN*_*pdn*_ in the *Nfkbia*^*-/-*^ settings to achieve fold differences between MOI 2 and 20 mirroring those determined for WT cells. Our modeling studies revealed that restoring the quantum of changes in *Virus*_*pdn*_ lessened the CHPV yield in *Nfkbia*^*-/-*^ cells at MOI 20 to those determined at MOI 2 (Fig 4e). Importantly, altering *Cell*_*death*_ also diminished substantially the progeny titer, whereas *IFN*_*pdn*_ had no impact. Therefore, a rise in the virus production rate and improved cell viability both appear to play a role in augmenting the CHPV yield with increasing input MOI in *Nfkbia*^*-/-*^ cells in the cell-saturating regime.

## Discussions

At early time points during pathogenesis, a limiting number of invading viruses present in the target tissue provides for sub-saturating low MOI cell infections. In the sub-saturating regime, viruses spread from infected to neighboring uninfected cells for propagation. Gradual accumulation of viruses in the infected tissue percolates into a transition to the cell-saturating high MOI regime at a later time point (Gutiérrez et al., 2010; Josefsson et al., 2011; Rüdiger et al., 2019). Combining experimental and mathematical studies, we determined the impact of input MOI on CHPV growth and antiviral cellular responses. As expected, increasing MOI boosted viral yield at sub-saturating low MOIs in our experiments, with type-1 IFNs playing a critical role in virus control (Fig 5). In the cell-suturing high MOI regime, where cell-to-cell viral transmission is negligible, cell death played a dominant role in restricting CHPV multiplication, leading to a paradoxical drop in viral yield upon increasing MOI (Fig 5). In a similar line, previous efforts to mathematically model Influenza virus growth indicated that limiting cellular resources and virus-induced apoptosis curtail viral production at high MOIs (Sidorenko et al., 2008; Yin & Redovich, 2018). Concordantly, Rüdiger et al. (2019) found that adjusting the apoptosis rate of Influenza virus-infected cells improves the model performance in the high MOI settings (Rüdiger et al., 2019). These mathematical studies also predicted that multiple infections of cells, typically achieved at cell-saturating MOIs, may undermine the antiviral state generated by type-1 IFNs (Rodriguez-Brenes et al., 2017a). Our present study reinforced this conceptual framework involving experimental evidence obtained using mutant cells to put forward a model where cell death curbs viral propagation mainly at high MOIs (Fig 5).

**Fig 5.**
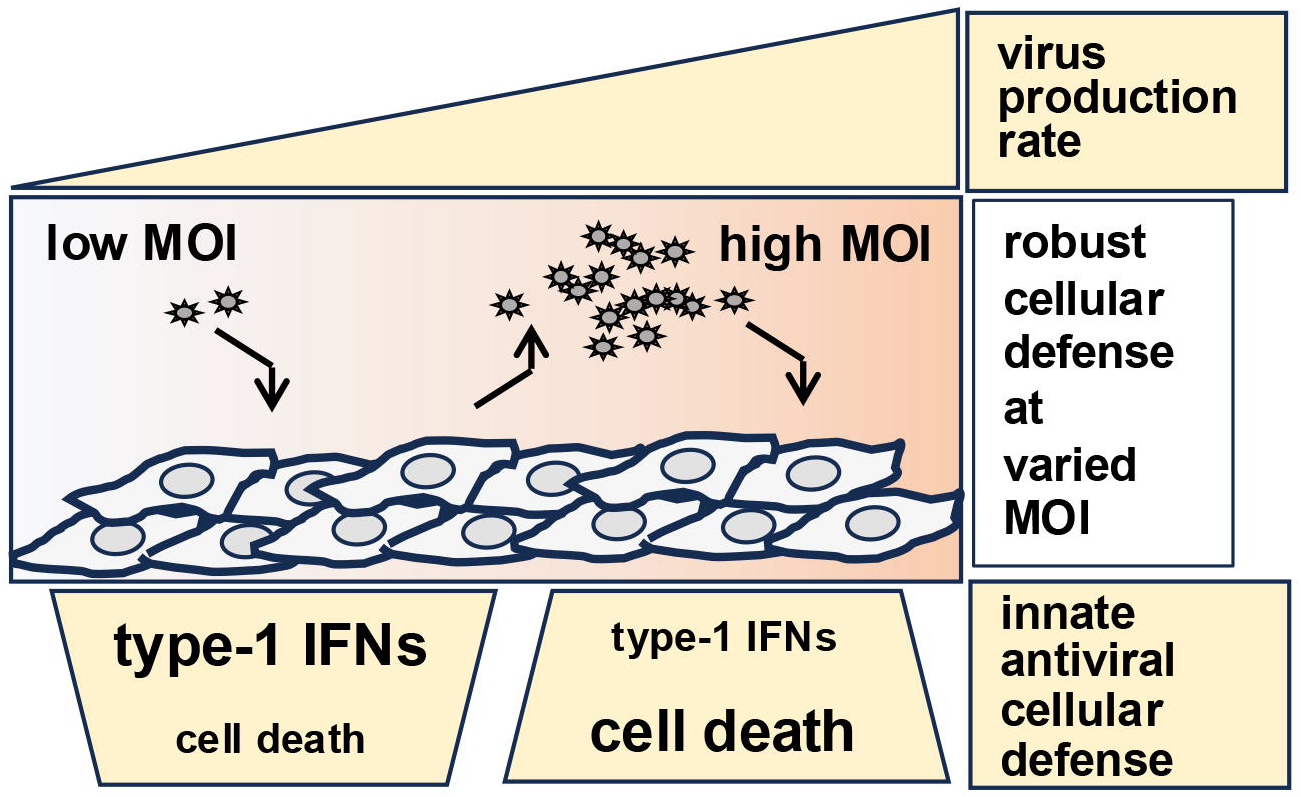
A model figure. A schema depicting the proposed collaboration between type-1 IFNs and cell death processes in limiting viral propagation at varied MOI.

Our investigation also captured multifaceted type-1 IFN mechanisms in the context of CHPV pathogenesis. Compared to MOI 2, CHPV infection of WT at MOI 20 led to an early cessation in IFNβ production that paralleled exacerbated cell death (Fig 1). Despite the impairment of type-1 IFN response, WT cells generated fewer progeny particles at MOI 20. Conversely, enhanced viability of *Nfkbia*^*-/-*^ cells prevented premature waning of IFNβ at MOI 20, and yet these knockout cells supported augmented yield upon raising MOI from 2 to 20 (Fig 4). These studies indicated that increased cell death at MOI 20 restricted not only viral multiplication but also type-1 IFN response. While emphasizing the mutually exclusive nature of antiviral cellular processes, our results strengthened the notion that type-1 IFNs are less consequential in regulating viral propagation at cell-saturating MOIs where cell death is pervasive. Taken together, we propose that distinct engagement of antiviral processes provides for robust cellular defense at varied MOIs.

We also scored a substantive effect of basal type-1 IFN signaling on CHPV growth that was confounded by the NF-κB pathway. It was previously reported that basal IFN signaling is required for boosting signal-induced IFN expressions (Ourthiague et al., 2015; Pantel et al., 2014; Yamane et al., 2019). We found that despite heightened virus multiplication, *Ifnar1*^*-/-*^ MEFs were less efficient at IFNβ production upon CHPV infection - a defect that could be attributed to the lack of basal IFN functions in these cells (Fig 3). Consistent with previous reports (Basagoudanavar et al., 2011), constitutive NF-κB signaling in *Nfkbia*^*-/-*^ MEFs promoted basally elevated IFNβ expressions (Fig 4). We speculate that this basal IFNβ level restricted the frequency of productive infection of *Nfkbia*^*-/-*^ cells, lessening CHPV multiplication and virus-induced type-1 IFN response in these mutants compared to WT MEFs. This IFN interference was partly overcome at very high MOIs (Rodriguez-Brenes et al., 2017b), presumably leading to an impactful increase in the virus production rate in *Nfkbia*^*-/-*^ cells upon raising MOI from 2 to 20. Future studies ought to combine flow cytometry-based experiments and mathematical analyses and employ compound knockout cells to more comprehensively dissect IFN and cell survival functions of NF-κB signaling in directing the cooperation between antiviral pathways. Similarly, the generalizability of the proposed model needs to be further explored in relation to other cytopathic viruses and physiologically relevant cell types.

While destroying the viral replicative niche, cell death also orchestrates danger-associated molecular pattern-mediated amplification of pathological inflammation in the infected tissue. Many of the anti-inflammatory drugs, including sodium salicylate and dexamethasone, commonly used for symptomatic management of viral diseases, suppress NF-κB signaling (Chamkouri et al., 2023; Di Bella et al., 2022; Yu et al., 2020). We anticipate that the newly described complex interplay between NF-κB functions and input MOIs during the progression of virus infection may bear broader significance for therapeutic intervention strategies in the future.

## Materials and Methods

### Primary cells and virus

Wild type (WT), *Ifnar1*^*-/-*^ and *Nfkbia*^*-/-*^ MEFs were generated from corresponding 12.5-13.5 day’s embryo and cultured in Dulbecco’s Modified Eagle Medium (DMEM) reconstituted with 10% BCS. Vero E6 cells were cultured in DMEM supplemented with 10% FBS. Chandipura virus (CHPV) strain 653514 was obtained from the National Institute of Virology, Pune and passaged in Vero E6 cells.

### Virus infection studies

Semi-confluent culture of primary MEFs were incubated with CHPV at varying MOI for 1.5hr in serum-free medium. Subsequent to virus adsorption, cells were washed with PBS and the culture was supplemented with DMEM containing 10% BCS. At various times post-infection, either culture supernatants were collected for measuring the titer of CHPV and the abundance of IFNβ or cells were harvested for assessing cell death.

### Virus plaque assay

Culture supernatants collected from infected cells were subjected to serial dilution in the incomplete media. Next, monolayers of Vero E6 cells were infected with these serial dilutions. After 1hr, inocula were removed, and Vero cells were dispensed with the overlay medium containing one part of DMEM and one part of 2% low melting agarose supplemented with 5% FBS at 4^0^C. Subsequently, cells were incubated at 37^0^C in a CO2 incubator for another 22hr and then fixed with 10% formaldehyde for 6hrs at room temperature. Viral plaques were visualized by staining the fixed VERO E6 monolayer with 0.2% crystal violet for 5 mins. Accordingly, viral titer in the culture supernatant was determined as PFU/ml.

### Cell death studies

CHPV-induced cell death was determined by crystal violet staining of adherent, live cells. Uninfected cells were used as controls. Cell death assays were performed using MEFs immortalized using NIH 3T3 protocol.

### Quantification of IFNβ levels

The abundance of IFNβ in the culture supernatant of infected MEFs was determined using a mouse IFNβ ELISA kit (PBL Assay Science, Piscataway, NJ) adhering to manufacturer’s protocol.

### Computational Modelling

To mathematically dissect CHPV growth at cell-saturating MOI, a three-state ordinary differential equations (ODE)-based model consisting of nine parameters were constructed (Fig 2a, Table 1). In this model, we assumed that all cells were infected at t=0 (*C(0)* =1). We considered that after a time delay of *t1*_*delay*_, infected cells (*C*) produced interferons (*IFN*) involving a rate constant *IFN*_*pdn*_ and progeny viruses (*V*) were generated involving a rate constant of *Virus*_*pdn*_. *IFN*_*inh*_ represented the rate constant for IFN-mediated inhibition of virus production by infected cells. After an additional delay of *t2*_*delay*_, cell death occurred involving a rate constant of *Cell*_*death*_. We considered that a minimum small fraction of cells, *C*_*basal*_, would remain alive even after prolonged infection. The degradation rate of IFNs and the rate for inactivation of viruses were represented using *IFN*_*deg*_ and *Virus*_*deg*_, respectively. Based on our experiments involving various knockout cells, we also considered a basal IFN level of 2.5 to 5.25 pg/ml in our model.

Briefly, the interferon concentration *IFN(t)* and the extracellular virus titer *V(t)* at a given time *t* are governed by the following ODEs:

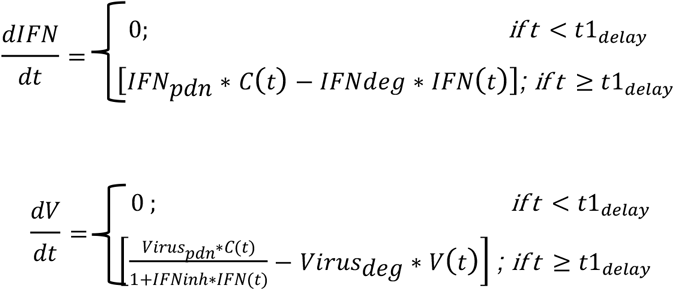

Similarly, the fraction of live cells *C(t)* at time *t* can be described by:

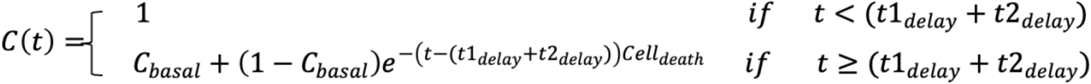

We conducted a comprehensive literature search to ascribe a range of initial values for these parameters for WT cells (Chakraborty et al., 2009; Timm & Yin, 2012). We then floated the parameter values. The final values were inferred by minimizing the combined least-square error between the model-predicted virus titers, interferon levels, and the fraction of live cells, and the observed experimental time course data for the MOI of 2 and 20 regimes, as described –

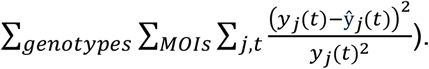

where *y*_*j*_(*t*) is the experimental measurement of variable j (*V(t), IFN(t), and C(t)*) at time t, and ŷ_*j*_(*t*) is the corresponding model prediction. This formula ensured that variables with larger absolute magnitudes (e.g., *V(t)*) did not dominate the fitting process, effectively balancing the contributions of *V(t), IFN(t), and C(t)*.

Parameters estimation was achieved by minimizing the pooled objective function using Excel’s GRG Nonlinear Solver. Accordingly, a set of parameter values were extracted for the WT settings and also for knockouts, adhering to following assumptions.

i. We noted that a rise in MOI from 2 to 20 in WT settings led to a less that 1.25 fold change in *t1*_*delay*_ from 2.5hr to 2hr; while *t2*_*delay*_ experienced an insignificant change from 3.25 to 3.24. Accordingly, we assumed that for the scope of our current analyses, *t1*_*delay*_ and *t2*_*delay*_ are insensitive MOIs. We then set the values of *t1*_*delay*_ and *t2*_*delay*_ in the WT settings at 2.5 and 3.5, respectively. Similar *t1*_*delay*_ and *t2*_*delay*_ values were fixed also for other genotypes.
ii. Because virus growth-inhibitory function and inactivation rate are intrinsic properties of IFNs, we assumed that *IFN*_*inh*_ and *IFN*_*deg*,_ are independent of the input MOI and cell genotypes. Accordingly, *IFN*_*inh*_ and *IFN*_*deg*_ values were kept identical between WT and *Nfkbia*^*-/-*^ systems. For *Ifnar*^*-/-*^ system, *IFN*_*inh*_ was set to zero for the want of functional type-1 IFN signaling, and *IFN*_*deg*,_ was kept similar to WT cells.
iii. Similarly, *Virus*_*deg*_ and *iv*) *C*_*basal*_ were persevered between genotypes.

### Statistical analysis

Error bars are presented as standard errors of the means (SEM) of three or more experimental replicates. Statistical significance was determined by Student’s t-test for computational studies and by two-way ANOVA followed by Tukey’s multiple comparison for experimental analyses.

## Acknowledgement

We sincerely thank present and past members of the Systems Immunology laboratory for help with experiments and constructive suggestions. We thank Mr Vijendra Kumar for technical help. Chandipura virus research in the PI’s laboratory is funded by the Department of Biotechnology, Govt. of India (BT/PR50989/COT/142/52/2023) and NII-Core. AS acknowledges the support of NIH-NIGMS via grant R35GM148351. Bhawna and SwB thank DST-INSPIRE and DBT, respectively, for research fellowships.

